# Know Your Alphabet: Conformational Noise, Latent-Space Encodings, and the Future of Structural Phylogenetics

**DOI:** 10.64898/2026.05.06.722973

**Authors:** Madeline Schmid, Yixiao Liu, Ashar J. Malik, David B. Ascher

**Author notes:** Correspondence to Ashar J. Malik, David B. Ascher.

## Abstract

Structural alphabets have transformed protein phylogenetics by enabling sequence-style alignment and maximum-likelihood inference to be applied directly to structural data. However, a coordinate-explicit alphabet, in which character states are derived from three-dimensional atomic positions, encodes not only evolutionary signal but also the conformational variability inherent to protein structure. This source of noise has not previously been quantified in a phylogenetic context, and no framework exists for comparing alphabets with respect to their conformational sensitivity. Here, we introduce the Normalised Noise Index (NNI), a Shannon entropy-based metric for quantifying conformational sensitivity in structural alphabet encodings, and apply it alongside ensemble-wide Robinson–Foulds (RF) variance as a framework for characterising the impact of conformational noise on phylogenetic inference. Across 3,749 single-chain NMR ensembles from the Protein Data Bank, we show that 3Di character variability is a pervasive feature of experimentally observed conformational spread, with NNI negatively correlated with within-ensemble structural stability. A 100 ns molecular dynamics simulation of myoglobin confirmed that thermal fluctuations alone are sufficient to generate comparable 3Di character variation and, in 2.9% of cases, to redirect maximum-likelihood tree search away from the expected topology in a 4-taxon globin benchmark with independently established relationships. Exhaustive enumeration of 4,800 conformational replicates across three NMR ensembles revealed that topological variance under 3Di encoding is approximately 1.7-fold greater than under structural distance, based on 11,517,600 pairwise RF comparisons, a source of uncertainty invisible to standard bootstrap analysis. By contrast, TEA, a sequence-derived structure-aware alphabet inferred from ESM-2 embeddings rather than directly from atomic coordinates, is insulated from conformational sampling by construction and yields zero topological variance across all conformational replicates, serving here as a noise-insulated reference rather than a proposed replacement for 3Di. Together, these results demonstrate that alphabet choice is a methodological variable in structural phylogenetics, and that the NNI metric and RF variance framework introduced here provide a practical basis for principled noise characterisation as new structural alphabets continue to emerge.

## Introduction

It is a long-standing principle of molecular evolution that protein structure is more conserved than primary sequence [1, 2, 3]. While the amino acid code can diverge into the “twilight zone” where homology is no longer detectable by sequence identity [4], the underlying three-dimensional fold often persists across geological timescales. Consequently, structural data has long been regarded as the superior substrate for uncovering deep evolutionary signals. Historically, however, the computational complexity of structural comparison limited these insights to small-scale analyses, while sequence-based phylogenetics flourished by utilizing mature algorithms for alignment and maximum-likelihood inference.

The introduction of the Foldseek 3Di (3D-interaction) alphabet [5, 6] represented a significant breakthrough in bridging this gap. By discretizing complex tertiary interactions into a string of structural characters, Foldseek enabled the field to adapt existing sequence-based phylogenetic workflows to structural data [6], building on earlier efforts to exploit structural conservation for deep-time inference [7, 8]. This enabled a paradigm in which structural alignments could be treated with the same mathematical rigor as their sequence counterparts; each character in an alignment column is interpreted as a discrete point on an evolutionary trajectory, contributing to the reconstructed history of a protein family. However, the validity of this interpretation depends on a property that has not previously been quantified at scale: whether structural alphabet characters are sufficiently stable across the conformational ensemble of a protein to serve as reliable evolutionary markers.

In the case of a coordinate-explicit alphabet like 3Di, however, this interpretation is complicated by a fundamental property of protein physics. The 3Di alphabet was designed to capture structural states with high fidelity for database searching, meaning it inherently captures the entirety of the structure, including the contributions of its dynamic nature. Unlike primary sequences, which are essentially static over the lifespan of an organism, protein structures are thermodynamic entities that explore ensemble properties through continuous thermal motion [3].

While a protein’s amino acid sequence remains constant, its structure “breathes,” exploring various conformational states that can result in several different 3Di representations for the same chain. This implies that 3Di-based representations frequently conflate evolutionary signal with conformational jitter. Without an additional analytical layer to filter out these thermodynamic contributions, the stochastic noise of molecular motion risks being misinterpreted as evolutionary divergence. Given the intricate relationship between geometric discretisation and the nature of structural change, deconvoluting signal from noise in coordinate-explicit representations remains an exceptionally difficult, and perhaps currently intractable, problem.

This ambiguity has a direct practical consequence that has not previously been quantified. A researcher building a structural phylogeny must select one deposited structure per protein, yet for NMR entries, dozens of equally valid conformational models exist, and crystal structures of the same protein in different ligand states or crystal forms are commonplace. If the 3Di encoding of a protein is sensitive to its conformational state, then the choice of which snapshot to use as input may itself influence the inferred topology. This represents a hidden source of uncertainty that is invisible to standard bootstrap analysis, which resamples alignment columns but cannot capture variance arising from conformational sampling.

As the challenges of explicit geometric representation have become clearer, the field of structural prediction has matured significantly. Modern protein language models have demonstrated that high-dimensional latent spaces can capture structural context directly from sequence data with remarkable power [9, 10, 11]. A recent advance in this area is TEA (The Embedded Alphabet) [12], a sequence-derived, structure-aware alphabet inferred from ESM-2 embeddings rather than directly from atomic coordinates. Because TEA anchors its structural representation to the invariant amino acid sequence via latent embeddings rather than fluctuating atomic positions, it is intrinsically insulated from conformational sampling of the same protein, making it a useful methodological reference for isolating the contribution of coordinate noise to phylogenetic variance.

As structural phylogenetics matures, it becomes increasingly important to define what a structural alphabet is actually measuring. In a coordinate-explicit alphabet such as 3Di, character states may reflect not only evolutionary divergence, but also the thermodynamic variability inherent to protein structure itself. To examine where structural signal begins to overlap with conformational noise, we evaluate the sensitivity of 3Di using two complementary settings. First, we analyse experimental NMR ensembles, which provide direct observations of conformational spread for a single underlying protein chain. Second, we use molecular dynamics (MD) simulations to test whether thermal fluctuations alone are sufficient to perturb 3Di encodings and alter downstream phylogenetic inference. Finally, we quantify the practical consequence of conformational sensitivity by exhaustively enumerating all possible conformational draws across three NMR ensembles, measuring both topological correctness and ensemble-wide Robinson–Foulds variance across 4,800 replicate trees. Throughout, we use the globin protein family as a controlled benchmark, chosen because its evolutionary relationships are unambiguously established across sequence [13], structure [14], and palaeontological evidence [15], providing a test system where the expected benchmark relationships are independently established. Together, these analyses provide the first direct estimate of the hidden topological uncertainty introduced by conformational alphabet choice.

By contrasting the coordinate-explicit representations with a latent-space alternative, we show that the choice of alphabet has direct consequences for how structural variation is interpreted in phylogenetic analysis. Structural alphabets remain a powerful advance for probing deep evolutionary relationships, but their phylogenetic utility depends on whether conformational variance is mistakenly treated as historical change. In this context, TEA serves as a noise-insulated methodological reference: because it is derived from sequence through latent embeddings rather than directly from fluctuating coordinates, it is intrinsically insulated from conformational sampling of the same protein, allowing the contribution of coordinate noise to phylogenetic variance to be isolated and measured. Together, these comparisons argue that future progress in structural phylogenetics will depend not only on improving structural representations, but also on a principled understanding of the noise characteristics each alphabet introduces; and that alphabet choice should be treated as a methodological variable, not an implementation detail.

## Methods

### Dataset Curation and Filtering

NMR ensembles were retrieved from a local mirror of the Protein Data Bank (updated February 2026). To isolate intramolecular conformational noise, we restricted the dataset to single-chain ensembles. Entries were required to exhibit model-wise length consistency and a minimum sequence length of 100 residues. This yielded a curated set of conformational replicates in which the underlying protein chain is constant while the 3D coordinates vary across experimentally observed models. Unless otherwise stated, the same model handling and filtering logic was applied to structures used in subsequent analyses. All structural encoding steps were performed using Foldseek version 8.ef4e960.

### 3Di Character Encoding

For each retained ensemble, all constituent models were extracted and converted into structural strings using the Foldseek 3Di alphabet. This process generated a set of model-specific 3Di strings of equal length for each PDB entry, providing a position-wise character representation of alternative conformational states for the same protein chain. Accordingly, where individual structures or selected structure sets were analysed in subsequent analyses, model-specific 3Di strings were generated and treated in the same manner unless otherwise noted.

### Quantifying Conformational Noise

Positional variability within 3Di ensembles was quantified via Shannon entropy. For a position *i*, with an observed frequency *p*_*i*_(*c*) for character *c* across *n* models, positional entropy was defined as:

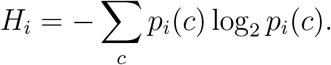

The ensemble-wide mean entropy 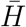 for a sequence of length *L* was calculated as:

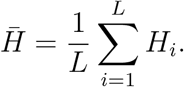

To standardise across ensembles of varying sizes, we computed a Normalised Noise Index (NNI) by dividing the mean entropy by the effective theoretical maximum, *H*_max_, defined as:

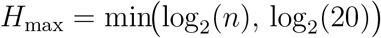

where *n* is the number of models and 20 is the size of the 3Di alphabet.

The Normalised Noise Index was then calculated as:

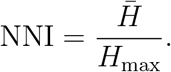

This cap ensures that the normalisation reflects the maximum entropy observable under both finite ensemble size and the 20-state capacity of the 3Di alphabet, and places the observed 3Di variability on a comparable scale bounded between 0 and 1.

### Structural Variance Analysis

To provide an empirical measure of conformational spread, all models within an ensemble were compared pairwise using Foldseek’s `alntmscore`. Self-comparisons were excluded. The structural stability of each ensemble was summarised by the mean pairwise TM-score:

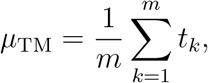

where *t*_*k*_ is the TM-score for the *k*th non-identical model pair and *m* is the total number of retained pairwise comparisons.

### Integration and Visualisation

The relationship between within-ensemble structural similarity (*µ*_TM_) and character-level noise (NNI) was assessed by merging the structural and entropic metrics. We visualised this relationship using a scatter plot to examine how within-ensemble structural stability relates to variability in the 3Di alphabet.

### Molecular Dynamics Simulation of Myoglobin

To generate dynamical conformational replicates, holo-myoglobin (protein chain and prosthetic heme) was isolated from solvent and non-heme hetero-components. Molecular dynamics (MD) simulations were performed in GROMACS [16] using the CHARMM36-feb2026 (LJ-PME/CGenFF 5.0) force field and TIP3P water model [17, 18]. The complex was centered in a cubic box with a 1.5 nm buffer, solvated, and neutralised with Na^+^ and Cl^−^ ions [19]. Following steepest descent minimisation (50,000 steps; 100 kJ mol^−1^ nm^−1^ tolerance), the system was heated under NVT conditions for 250 ps using simulated annealing (50 to 310 K) and V-rescale temperature coupling with a 2 fs timestep [19]. Subsequent NPT equilibration was conducted for 500 ps at 310 K and 1 bar using the isotropic c-rescale barostat [19]. A 100 ns production simulation was executed under NPT conditions (310 K, 1 bar) using V-rescale/c-rescale coupling, PME electrostatics, and LJ-PME van der Waals treatment [19]. Position restraints were maintained on the protein and heme during equilibration and removed for production [19]. Trajectory coordinates were recorded every 10 ps, yielding 10,000 candidate conformational snapshots for downstream analysis [19]. All simulations were performed using GROMACS 2026.1.

### Trajectory Analysis and Noise Quantification

To obtain an approximately independent set of sampled conformations, the statistical inefficiency (*g*) of the production trajectory was estimated using the Flyvbjerg– Petersen block averaging algorithm applied to the backbone RMSD series. Snapshots (*n ≈* 69) were then extracted at intervals of *g* to serve as representative conformational replicates. To assess the impact of thermal motion on the 3Di alphabet, these snapshots were converted to 3Di sequences, and the ensemble-wide NNI was calculated using the positional Shannon entropy framework described above. Structural similarity across the trajectory was quantified by computing the mean pairwise TM-score (*µ*_TM_) for all extracted frames using Foldseek. This dual-track analysis enabled direct comparison between character-level noise in the MD-derived ensemble and the conformational spread observed across experimental NMR ensembles.

### Preparation of Static 3Di Reference Sequences

To establish the evolutionary background for the quartet benchmark, static reference structures for *Homo sapiens* haemoglobin*−α* (`4HHB_A`), *Homo sapiens* haemoglobin*−β* (`4HHB_B`), and *Physeter macrocephalus* myoglobin (`1MYF_A`) were isolated from their respective asymmetric units. These were combined with MD-derived conformational replicates of *Sus scrofa* myoglobin (`1MYG A`) to form the 4-taxon benchmark. These were converted into 3Di structural sequences using Foldseek. During conversion, histidine protonation states were standardised to canonical 3-letter codes to ensure nomenclature consistency between the experimental crystal structures and the MD-derived replicates.

### Construction of the Evolutionary Perturbation Benchmark

A benchmark quartet was assembled from the four static reference sequences to define the expected topology. To quantify the impact of conformational noise, replicate datasets were generated by iteratively substituting the static `1MYG A` sequence with each uncorrelated MD-derived myoglobin conformer, while maintaining the other three taxa in their static states. This controlled substitution framework ensured that any observed topological variance arose from the structural fluctuations sampled during molecular dynamics simulation of a single target taxon.

### Substitution Model Parameterisation

Since both 3Di and TEA were originally distributed with substitution matrices for structural alignment, we derived corresponding PAML-format exchangeability matrices to enable maximum-likelihood inference. Following the same procedure used for the Foldseek 3Di alphabet [6], the original substitution scores were transformed into exchangeabilities using the background state frequencies *f*_*j*_ and the matrix-specific scaling factor *λ* provided with each matrix:

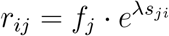

The resulting values were written as 20-state lower-triangular matrices together with their associated equilibrium frequencies, following the format expected by IQ-TREE. This transformation allowed both structural alphabets to be treated as discrete-state substitution models during phylogenetic inference, such that tree search was guided by alphabet-specific structural substitution patterns rather than standard amino acid models.

### Alignment, Inference, and Topology Scoring

Structural sequences were encoded using Foldseek version 8.ef4e960 for 3Di and the `tea convert` utility for TEA. Encoded sequences were aligned using ClustalW2 version 2.1 [20] with the author-provided substitution matrices corresponding to each alphabet. Phylogenetic trees were then inferred using IQ-TREE version 1.6.12 [21] under a maximum-likelihood framework, with tree searches guided by the corresponding custom-derived exchangeability matrices rather than standard amino acid models. Taxon labels were normalised to allow for consistent scoring of recovery of the expected bipartition:

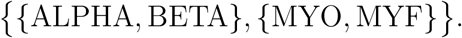

The inferred topologies were parsed using bipartition-matching to determine the proportion of trees supporting the expected quartet relationship, thereby quantifying the robustness of the alphabet to conformational noise.

### Exhaustive NMR Replicate Benchmark and Topological Stability Analysis

To assess hidden topological instability in a larger conformational benchmark, we constructed an 8-taxon dataset comprising five static crystal structures and three NMR-derived sources sampled across all available models. Static taxa included the *α* and *β* chains of *Anser indicus* haemoglobin (`1HV4_A`, `1HV4_B`), the *α* and *β* chains of *Danio rerio* haemoglobin (`1T1N_A`, `1T1N_B`), and *Sus scrofa* myoglobin (`1MYG_A`). Variable taxa were represented by all models from *Homo sapiens* haemoglobin*−α* (`2H35_A`, *n* = 20), *Homo sapiens* haemoglobin*−β* (`2H35_B`, *n* = 20), and *Physeter macrocephalus* myoglobin (`1MYF_A`, *n* = 12). All possible combinations of one model from each variable source were enumerated, yielding 4,800 replicate datasets.

### Replicate Tree Construction Under Distance and 3Di Frameworks

For each replicate, trees were inferred under two alternative frameworks. In the 3Di framework, replicate structures were converted to Foldseek 3Di strings, aligned with ClustalW2 using the 3Di substitution matrix, and analysed with IQ-TREE under the corresponding custom-derived exchangeability matrix described above. In the distance framework, all replicate structures were compared all-against-all using Foldseek, and pairwise `alntmscore` values were converted to distances as

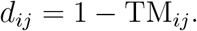

These distances were assembled into replicate-specific distance matrices, from which neighbour-joining trees were inferred using `scikit-bio`. For both frameworks, model-specific labels were collapsed back to source-level taxon identifiers prior to downstream scoring. Neighbour-joining trees were constructed using scikit-bio version 0.6.3 under Python 3.11.12. The random seed for combination enumeration was fixed at 42 to ensure reproducibility of replicate ordering.

### Group Recovery and Ensemble-Wide RF Analysis

Each replicate tree was assessed for recovery of the expected *α, β*, and *myoglobin* groupings. For each grouping, monophyly was evaluated by identifying the least common ancestor of the expected taxa and testing whether that clade contained only the members of the target set. In addition, a clade purity metric was calculated as the proportion of taxa within the least common ancestor that belonged to the expected group. An overall recovery flag was assigned when all expected groupings were simultaneously recovered within a replicate tree.

To quantify topological stability across the full replicate ensemble, all trees inferred under a given framework were compared pairwise using normalised Robinson– Foulds (RF) distance. Bipartitions were extracted from midpoint-rooted trees, and the normalised RF distance between two trees was computed as the size of the symmetric difference between their bipartition sets divided by the maximum possible difference for that pair. The resulting RF distribution was summarised by its mean, median, and standard deviation, providing an ensemble-level measure of hidden topological variability under each inference framework.

As TEA encodes amino acid sequences rather than conformational state, all NMR models of the same protein produce identical TEA characters; accordingly, TEA trees are invariant to conformational sampling and the RF distance between any two TEA replicates is zero by construction.

### Web Application Availability

Part of the analysis pipeline described above has been made available as a web application at https://biosig.lab.uq.edu.au/noisyalphabet. The application was implemented using the Flask web framework (Python), served via Nginx, and containerised using Docker as part of the Structome suite [22, 23]. The frontend was built with HTML, CSS, and JavaScript, incorporating the Mol* molecular viewer [24] for interactive structure visualisation. Users may upload any single-chain NMR ensemble in PDB format and receive the NNI, mean pairwise TM-score, per-column entropy statistics, and placement on the PDB-wide conformational noise landscape described in this work.

## Results

### Conformational noise in the 3Di alphabet across NMR ensembles

To characterise the relationship between structural stability and 3Di conformational noise at scale, we analysed 3,749 single-chain NMR ensembles retrieved from the Protein Data Bank, each comprising at least two models and a minimum of 100 residues. For each ensemble, the mean pairwise TM-score was computed as a measure of within-ensemble structural stability, and the Normalised Noise Index (NNI) was computed from the column-wise Shannon entropy of the 3Di character alignment across all constituent models.

Across the dataset, NNI values ranged from near zero to 0.58, with a mean of 0.17, indicating that conformational noise in the 3Di encoding is a pervasive rather than exceptional feature of NMR ensembles. A negative relationship between mean TM-score and NNI was observed across the full dataset (Figure 1): ensembles with lower mean pairwise TM-score, reflecting greater within-ensemble structural spread, consistently exhibited higher 3Di character variability. Notably, this relationship was not restricted to structurally unstable outliers; ensembles with mean TM-scores approaching 1.0, indicating very tight conformational distributions, still exhibited non-zero NNI values, demonstrating that even minor thermal fluctuations are sufficient to alter 3Di character assignments at a subset of positions.

**Figure 1:**
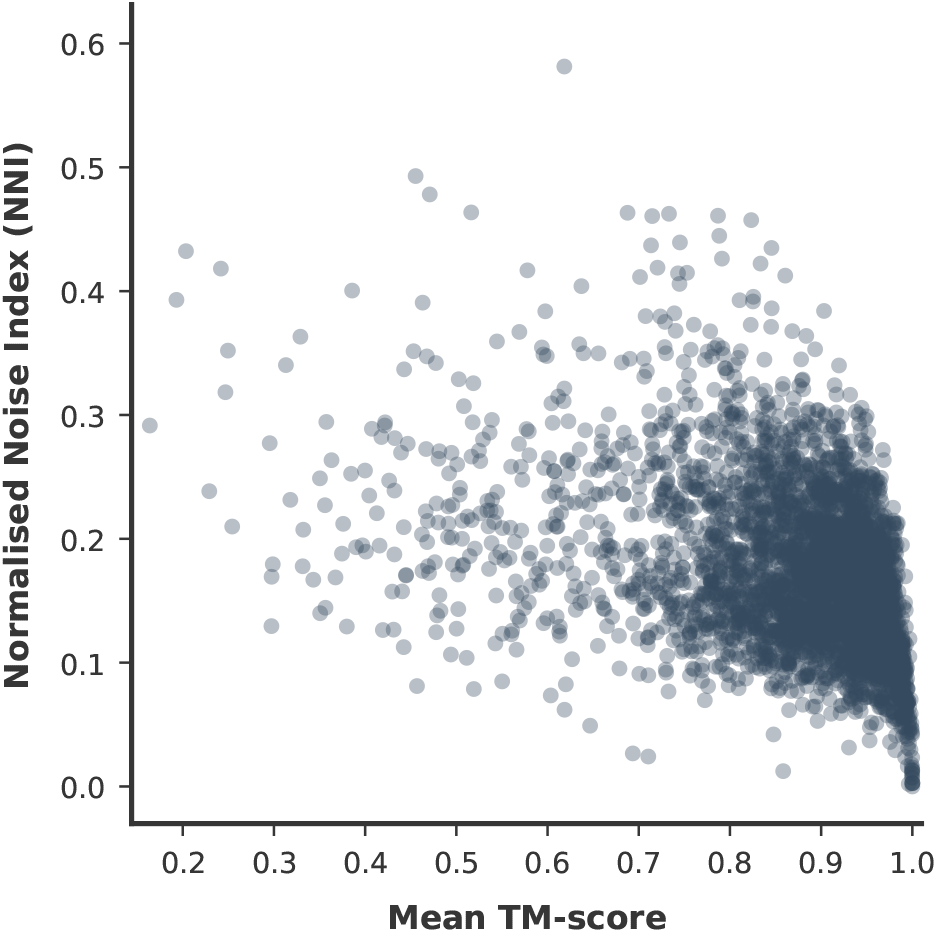
Relationship between within-ensemble structural stability and 3Di conformational noise across 3,749 single-chain NMR ensembles from the Protein Data Bank. Each point represents one ensemble. The x-axis shows the mean pairwise TM-score, a measure of structural stability across models. The y-axis shows the Normalised Noise Index (NNI), quantifying the mean column-wise Shannon entropy of the 3Di character alignment normalised by the theoretical maximum for that ensemble size, capped at the 20-state 3Di alphabet ceiling. A negative relationship is observed across the full dataset, with structurally unstable ensembles encoding greater conformational noise in the 3Di alphabet. Non-zero NNI values persist even among structurally tight ensembles (mean TM-score > 0.95), indicating that minor thermal fluctuations are sufficient to alter 3Di character assignments. All 3,749 ensembles are shown across the full range of within-ensemble structural stability, including intrinsically disordered proteins and engineered mutants at the lower end of the TM-score distribution.

### Molecular dynamics simulation reveals conformational sensitivity of 3Di topology

A 100 ns molecular dynamics simulation of myoglobin (`1MYG A`) yielded 69 statistically independent conformational snapshots following Flyvbjerg–Petersen subsampling of the backbone RMSD series (Figure S1). The MD ensemble exhibited a mean pairwise TM-score of 0.963 and a Normalised Noise Index of 0.126, placing it within the distribution of experimentally observed NMR ensembles shown in Figure 1 and confirming that thermal fluctuations can generate measurable 3Di character variation at a level comparable to solution NMR conformational spread.

For four taxa there are exactly three possible unrooted tree topologies (Figure 2). The expected benchmark topology groups the two haemoglobin chains together and the two myoglobins together — (ALPHA + BETA) — (MYO + MYF) — consistent with the known evolutionary relationships within the globin family. Each of the 69 MD-derived conformational snapshots was substituted in turn as the respective myoglobin replicate in a 4-taxon dataset alongside static reference sequences for human haemoglobin*−α* (`4HHB_A`), human haemoglobin*−β* (`4HHB_B`), and sperm whale myoglobin (`1MYF_A`). Together with the static PDB reference structure, these 69 MD-derived conformers yielded 70 total quartet realisations for phylogenetic analysis. Under 3Di encoding, 68 of 70 quartet realisations (97.1%) recovered the expected topology (a), while 2 realisations (2.9%) recovered topology (b), in which pig myoglobin grouped with a haemoglobin chain rather than with sperm whale myoglobin. Topology (c) was never recovered. Under TEA encoding, the single quartet tree recovered the expected topology (a). Critically, each replicate tree was constructed from a genuinely different conformational snapshot rather than a resampled alignment, hence this is not to be interpreted as a bootstrap analysis. The two topologically discordant 3Di trees therefore represent cases in which thermal conformational noise in the 3Di encoding was sufficient to redirect the maximum-likelihood tree search toward an alternative evolutionary grouping.

**Figure 2:**
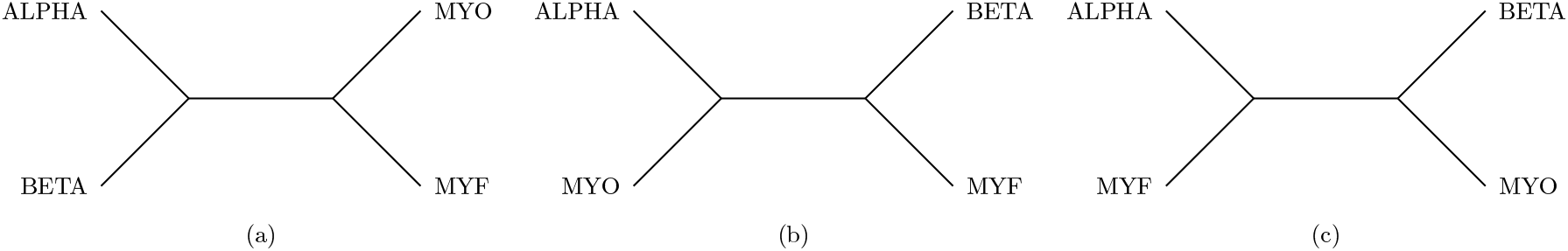
The three possible unrooted topologies for the 4-taxon globin benchmark comprising human haemoglobin*−α* (ALPHA, `4HHB_A`), human haemoglobin*−β* (BETA, `4HHB_B`), sperm whale myoglobin (MYF, `1MYF_A`), and pig myoglobin (MYO, `1MYG_A`). Topology (a) represents the expected benchmark grouping, placing haemoglobin chains together and myoglobins together. Under 3Di encoding, 68 of 70 quartet realisations (97.1%) recovered topology (a), while 2 realisations (2.9%) recovered topology (b). Topology (c) was never recovered. Under TEA encoding, the single quartet tree recovered topology (a). Each replicate represents a genuinely distinct conformational snapshot rather than a resampled alignment.

### Exhaustive conformational sampling reveals hidden topological variance in 3Di-based inference

To quantify the hidden topological uncertainty introduced by conformational alphabet choice, we exhaustively enumerated all possible combinations of NMR models across three variable sources, `2H35_A` (20 models), `2H35_B` (20 models), and `1MYF_A` (12 models), combined with five static crystal structure chains (`1HV4_A`, `1HV4_B`, `1T1N_A`, `1T1N_B`, `1MYG_A`), yielding 4,800 replicate 8-taxon datasets. For each replicate, trees were inferred independently under two frameworks: 3Di maximum-likelihood and TM-score distance neighbour-joining. Group recovery was then assessed for three expected monophyletic groupings: *α*-globins, *β*-globins, and myoglobins.

Under the distance framework, all three groupings were simultaneously recovered in 2,996 of 4,800 replicates (62.4%). Under 3Di encoding, simultaneous recovery dropped to 1,454 of 4,800 replicates (30.3%), representing a 2.1-fold reduction in simultaneous recovery of the expected groupings.

Per-group recovery rates revealed that the overall 3Di underperformance was driven entirely by the *β−*haemoglobin chain: *β−*globin recovery fell from 98.5% (distance) to 30.4% (3Di), while *α−*globin recovery was actually higher under 3Di than distance (99.8% vs 63.8%), and myoglobin grouping was robust under both frameworks (99.9% vs 100.0%). Full per-group and overall recovery statistics are summarised in Table 1. This asymmetry indicates that conformational noise in 3Di encoding is not uniformly distributed across globin subtypes, and that the *β−*haemoglobin chain is disproportionately sensitive to the conformational state of the human haemoglobin NMR ensemble used here.

**Table 1:**
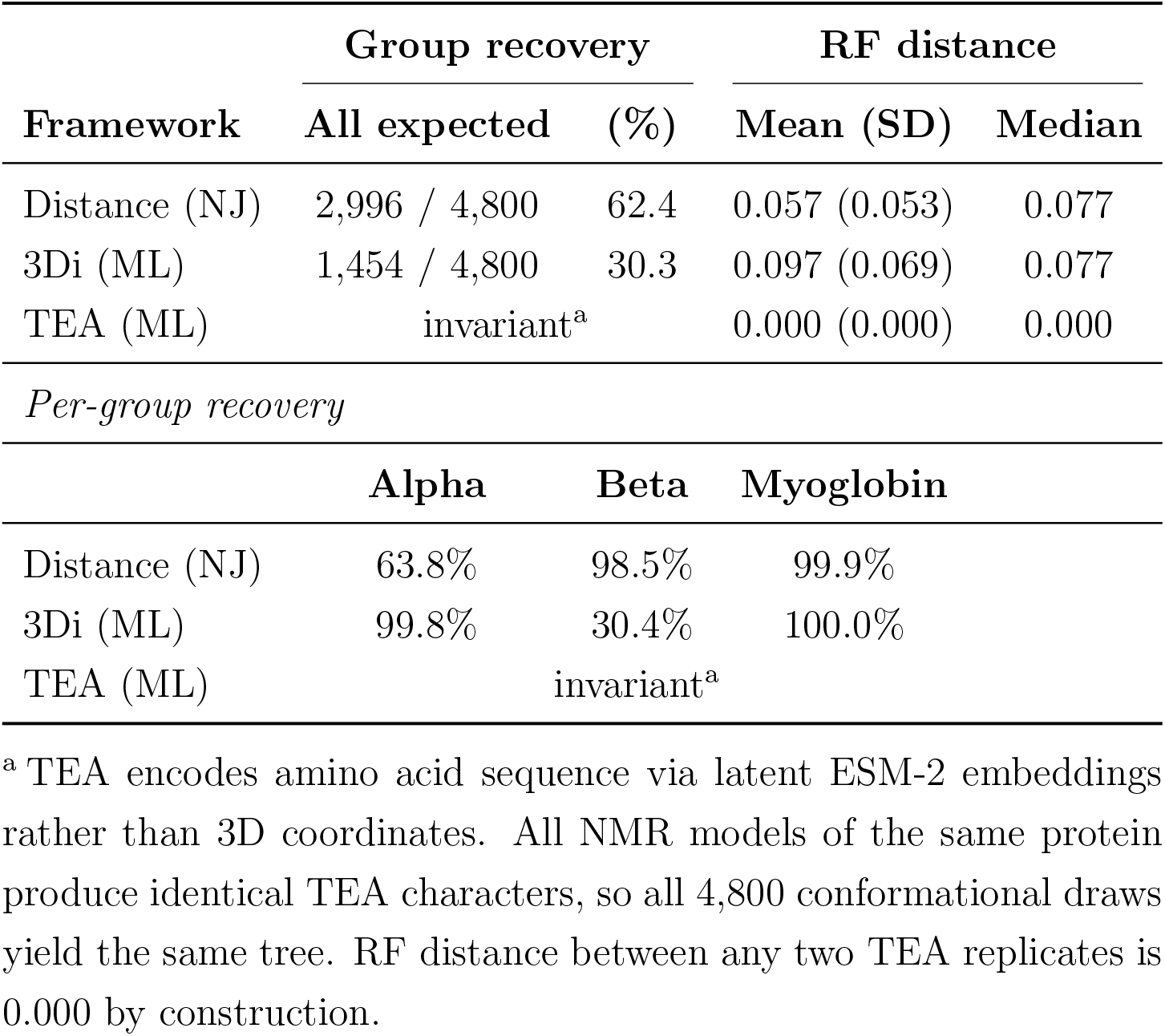
Topological recovery and ensemble-wide RF variance across 4,800 exhaustive NMR conformational replicates under three inference frameworks. Group recovery reports the number and percentage of replicates in which the expected monophyletic grouping was recovered. RF statistics summarise the full pairwise RF distance distribution across all 11,517,600 replicate pairs. TEA RF is zero by construction as TEA encodes amino acid sequence rather than conformation.

| Framework | Group recovery |  | RF distance |  |
| --- | --- | --- | --- | --- |
|  | All expected | (%) | Mean (SD) | Median |
| Distance (NJ) | 2,996 / 4,800 | 62.4 | 0.057 (0.053) | 0.077 |
| 3Di (ML) | 1,454 / 4,800 | 30.3 | 0.097 (0.069) | 0.077 |
| TEA (ML) | invariant<sup>a</sup> |  | 0.000 (0.000) | 0.000 |
| *Per-group recovery* |  |  |  |  |
|  | Alpha | Beta | Myoglobin |  |
| Distance (NJ) | 63.8% | 98.5% | 99.9% |  |
| 3Di (ML) | 99.8% | 30.4% | 100.0% |  |
| TEA (ML) | invariant<sup>a</sup> |  |  |  |

<sup>a</sup> TEA encodes amino acid sequence via latent ESM-2 embeddings rather than 3D coordinates. All NMR models of the same protein produce identical TEA characters, so all 4,800 conformational draws yield the same tree. RF distance between any two TEA replicates is 0.000 by construction.

Critically, the primary purpose of this analysis was not to assess which framework recovered the expected groupings more often, but to quantify how much the inferred topology varies depending on which conformational snapshot was selected as input, a source of uncertainty in structural phylogenetics. To measure this ensemble-wide variance, all 11,517,600 pairwise normalised Robinson–Foulds (RF) distances were computed across the full 4,800-tree ensemble for each framework. Under 3Di encoding, any two trees drawn from different conformational combinations differed by a mean normalised RF of 0.097 (median 0.077, std 0.069). Under the distance framework this variance was approximately 1.7-fold lower, with a mean normalised RF of 0.057 (median 0.077, std 0.053). The identical median values reflect the discrete quantisation of the 8-taxon RF space, in which only a small number of distinct bipartition configurations are possible; the mean and standard deviation are therefore the more discriminating statistics for comparing the two distributions. Under TEA encoding, the RF distance between any two replicates would be zero by construction. Because TEA encodes amino acid sequence rather than conformation, all 4,800 conformational draws would produce identical TEA characters for each taxon, and the same TEA tree would therefore be obtained for every possible combination.

## Discussion

The Foldseek 3Di alphabet was designed for ultrafast structural database searching, not for phylogenetic inference. Its adaptation to phylogenetics is a natural and intellectually compelling extension that opens genuinely new analytical territory, enabling sequence-style alignment and maximum-likelihood inference to be applied directly to structural data at scale. However, tools adapted beyond their original design scope carry implicit assumptions that require scrutiny. The foundational assumption of character-based phylogenetics is that alignment columns represent stable, heritable evolutionary states. For amino acid sequences this assumption is well justified; the primary sequence of a protein is invariant across the conformational ensemble explored during its lifetime. For a coordinate-explicit structural alphabet, where character states are derived from three-dimensional atomic positions, this assumption is not guaranteed. The present work does not argue that 3Di-based structural phylogenetics is without merit; instead it argues that conformational sensitivity is a property of coordinate-explicit encoding that deserves explicit attention, and that its consequences for downstream inference can be measured and compared across alphabets.

The field of structural alphabet-based phylogenetics is still at an early and rapidly evolving stage. Until recently, 3Di represented the only practical option for character-based structural phylogenetic inference. The emergence of TEA as an alternative representation, derived from sequence through protein language model embeddings rather than from atomic coordinates, yet still encoding structure-aware characters, expands the available repertoire and creates an opportunity for principled methodological comparison. When two alphabets with fundamentally different encoding philosophies become available, it becomes possible to ask not only whether structural phylogenetics works, but which properties of a structural encoding are beneficial and which introduce unwanted variance. The present work takes advantage of this opportunity. While the broader phylogenetic utility of TEA remains to be established through large-scale validation, the specific source of uncertainty exposed here, conformational sensitivity in coordinate-explicit encoding, is one that TEA is, by design, insulated from. This does not make TEA universally superior; rather, it makes TEA a useful reference point for isolating the contribution of conformational noise to phylogenetic variance, and a proof of principle that structure-aware alphabets need not be directly coupled to thermodynamic fluctuations in atomic coordinates.

The results presented here demonstrate that this sensitivity is both real and consequential. Across 3,749 single-chain NMR ensembles, 3Di character variability was a pervasive feature of the dataset rather than an exceptional one, with non-zero NNI values persisting even among structurally tight ensembles. A 100 ns molecular dynamics simulation of *Sus scrofa* myoglobin confirmed that thermal fluctuations alone, without any evolutionary divergence, are sufficient to generate 3Di character variation at a level comparable to experimental NMR conformational spread, and in 2.9% of cases sufficient to redirect maximum-likelihood tree search away from the expected topology in the 4-taxon globin benchmark. The exhaustive NMR replicate analysis extended this finding to a larger combinatorial space, revealing that the choice of which conformational snapshot to use as input introduces a 1.7-fold greater topological variance under 3Di encoding than under structural distance, as measured by pairwise RF distance across 11,517,600 replicate tree comparisons. This variance is invisible to standard bootstrap analysis, which resamples alignment columns but cannot capture uncertainty arising from conformational sampling. The globin family was deliberately chosen as the test system for this analysis because its evolutionary relationships are unambiguous and independently established across sequence, structure, and palaeontological evidence, making it an appropriate and reusable benchmark for evaluating structural phylogenetic methods where the expected relationships are independently established.

TEA escapes the conformational noise problem by construction. Because it derives its structural characters from amino acid sequences via ESM-2 latent embeddings rather than from fluctuating atomic coordinates, it is intrinsically insulated from conformational sampling of the same protein. This property made it a useful methodological control in the present work, not as a claim that TEA is the correct or final solution for structural phylogenetics, but as a demonstration that conformationally invariant structural encodings are achievable and produce measurably different noise profiles. TEA has not been validated at scale for phylogenetic applications and, furthermore, inherits the assumptions and limitations of transformer-based protein language models, which introduce their own sources of uncertainty. Of particular relevance is the observation that the same ESM-2 embedding space exploited by TEA for alphabet generation is now being actively investigated in parallel work for direct structural ensemble sampling [25, 26]; meaning the boundary between sequence-derived and coordinate-derived structural representations is itself in flux. As this area matures, the noise characteristics of embedding-derived alphabets will themselves require principled evaluation using frameworks such as the one introduced here.

The Normalised Noise Index and the pairwise RF variance approach introduced in this work are alphabet-agnostic. They can be applied to any structural encoding, current or future, to characterise how sensitive the resulting phylogenetic inference is to conformational sampling. As the field develops new structural alphabets, we argue that noise characterisation of this kind should be a standard part of the validation pipeline rather than an afterthought. The globin benchmark introduced here, with its independently verified phylogeny, multiple NMR ensembles of known conformational spread, and exhaustive combinatorial replicate design, provides a reusable test case for this purpose.

The scope of this work is deliberately focused. We chose one protein family in a controlled benchmark setting where the expected relationships are independently established. This is sufficient to establish the principle and introduce the quantitative framework, but it is not sufficient to make generalised claims about the performance of either alphabet across the full diversity of structural phylogenetics applications. The practical implication is specific and bounded: when 3Di or any coordinate-explicit structural alphabet is used for phylogenetic inference, the choice of input structure is a methodological variable that can influence the inferred topology, and this source of uncertainty should be acknowledged and where possible quantified alongside standard measures of bootstrap support. Because the 3Di and distance frameworks differ in both representation and inference procedure, this comparison should be interpreted as an end-to-end comparison of practical workflows rather than as an isolation of alphabet effects alone.

### Disclaimer

The comparison between 3Di and TEA presented in this work should be interpreted with care. The two alphabets differ not only in their source of signal, coordinate-explicit versus sequence-derived, but also in their underlying substitution matrices, scoring philosophies, and the assumptions embedded in their respective alignment frameworks. These differences mean that observations about alignment conservation, gap frequency, or column entropy cannot be directly compared between alphabets as independent lines of evidence for one encoding’s superiority over another, since such statistics are partly a function of matrix design rather than purely of biological signal. The RF variance result is the most robust comparative metric presented here precisely because it measures topological consistency across conformational replicates rather than alignment quality, and is therefore not confounded by matrix design differences. All comparisons in this work should be understood as illustrative of a principle: that coordinate-explicit and sequence-derived structural alphabets have fundamentally different noise characteristics, rather than as a comprehensive performance evaluation of either tool.

## Conclusion

The results presented here establish that conformational noise is a measurable and consequential property of coordinate-explicit structural alphabets, and that its impact on phylogenetic inference can be quantified using the framework introduced in this work. Across 3,749 NMR ensembles, a 100 ns molecular dynamics trajectory, and 4,800 exhaustive conformational replicates, we show that the choice of input structure is not a neutral decision when using 3Di for phylogenetic inference. Instead, it is a methodological variable that can introduce topological variance. TEA, as a latent-space alternative anchored to the invariant amino acid sequence, is insulated from this source of noise by construction and therefore provides a proof of principle that structure-aware alphabets need not inherit thermodynamic uncertainty from explicit atomic coordinates. As structural phylogenetics continues to mature and new alphabets enter the field, we argue that principled characterisation of conformational noise should become a standard part of alphabet evaluation, and that the NNI metric, the RF variance framework, and the globin benchmark introduced here provide a practical starting point for that characterisation.

## Data and Web Application Availability

The PDB-wide conformational noise landscape described in Figure 1 is interactively accessible via the NoisyAlphabet web application at https://biosig.lab.uq.edu.au/noisyalphabet. Users may upload any single-chain NMR ensemble in PDB format and receive the Normalised Noise Index, mean pairwise TM-score, and per-column 3Di entropy statistics for their structure of interest, together with its placement on the PDB-wide scatter plot. This provides a direct means of reproducing and contextualising the Figure 1 analysis for any NMR ensemble of interest without requiring local installation of any software.

## Use of Artificial Intelligence

Artificial intelligence tools were used to assist in improving the readability of this manuscript. All scientific content, data analysis, interpretation, and conclusions are the sole responsibility of the authors.

## Acknowledgements

D.B.A. is supported by an NHMRC Investigator Grant (GRNT2041888). This research was supported by the NVIDIA Academic Grant Program.

