## Supplementary material for "Know Your Alphabet: Conformational Noise, Latent-Space Encodings, and the Future of Structural Phylogenetics": Figure S1

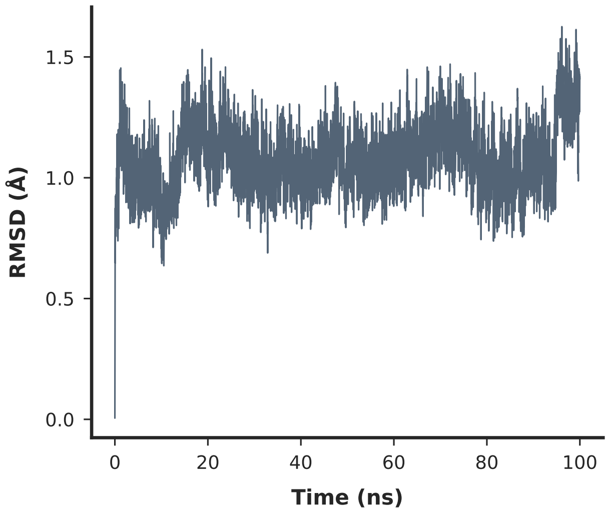


Figure S1: Backbone RMSD over the 100 ns production molecular dynamics trajectory of *Sus scrofa* myoglobin (`1MYG_A`). RMSD was computed relative to the minimised starting structure using backbone heavy atoms. The trajectory exhibits stable behaviour after initial equilibration, confirming that the simulation reached a stable sampling regime prior to frame extraction. The statistical inefficiency $g$ estimated by the Flyvbjerg–Petersen block averaging algorithm applied to this series determined the frame extraction interval, yielding approximately 69 statistically independent conformational snapshots for downstream analysis.
